# Challenges in estimating heritability of phase polyphenism: insights from measured and simulated data in the desert locust

**DOI:** 10.1101/149542

**Authors:** Hélène Jourdan-Pineau, Benjamin Pélissié, Elodie Chapuis, Floriane Chardonnet, Christine Pagès, Antoine Foucart, Laurence Blondin, Cyril Piou, Marie-Pierre Chapuis

## Abstract

Quantitative genetics experiments aim at understanding and predicting the evolution of phenotypic traits. Running such experiments often bring the same questions: Should I bother with maternal effects? Could I estimate those effects? What is the best crossing scheme to obtain reliable estimates? Can I use molecular markers to spare time in the complex task of keeping track of the experimental pedigree?

We explored those practical issues in the desert locust, *Schistocerca gregaria* using morphologic and coloration traits, known to be influenced by maternal effects. We ran quantitative genetic analyses with an experimental dataset and used simulations to explore i) the efficiency of animal models to accurately estimate both heritability and maternal effects, ii) the influence of crossing schemes on the precision of estimates and iii) the performance of a marker-based method compared to the pedigree-based method.

The simulations indicated that maternal effects deeply affect heritability estimates and very large datasets are required to properly distinguish and estimate maternal effects and heritabilities. In particular, ignoring maternal effects in the animal model resulted in overestimation of heritabilities and a high rate of false positives whereas models specifying maternal variance suffer from lack of power. Maternal effects can be estimated more precisely than heritabilities but with low power. To obtain better estimates, bigger datasets are required and, in the presence of maternal effects, increasing the number of families over the number of offspring per families is recommended. Our simulations also showed that, in the desert locust, using relatedness based on available microsatellite markers may allow reasonably reliable estimates while rearing locusts in group.

In the light of the simulation results, our experimental dataset suggested that maternal effects affected various phase traits. However the statistical limitations, revealed by the simulation approach, didn’t allow precise variance estimates. We stressed out that doing simulations is a useful step to design an experiment in quantitative genetics and interpret the outputs of the statistical models.

## Introduction

Trait evolution directly depends on the phenotypic variation transmitted across generations by genetic inheritance, parental effect or even cultural and ecological inheritance (Danchin, Charmantier, Champagne, *et al.*, 2011). Therefore, predicting the evolutionary potential of a phenotypic trait requires quantifying the amount of phenotypic variation due to genetic, maternal (or more generally parental) and environmental effects, which is the general objective of quantitative genetics (Lynch & Walsh, 1998). Quantitative genetics experiments rely on the phenotypic resemblance of related individuals and are therefore based on controlled crossings and phenotypic measurements of individuals of known pedigree. Running a quantitative genetics experiment for the first time on a new model species can be challenging and requires a careful consideration of the crossing scheme, pedigree inference and statistical model.

First, heritability is estimated by measuring phenotypes of individuals of known degrees of relatedness. To obtain such data, it is necessary to use a population with a pedigree data ranging over several generations or to design an experiment with specific relatedness classes. Thus, in the laboratory, controlled crosses are required and the chosen crossing scheme has a real impact on the nature and precision of the estimates. For example, full-sib design only gives an estimate of the broad-sense heritability (H^2^) that contains all the genetic variance in the form of additive, dominance and epistatic allele effects (divided by the phenotypic variance) whereas a half-sib/full-sib design gives an estimate of the narrow-sense heritability (h^2^) containing only the additive effect of the genetic variance (Lynch & Walsh, 1998). Since response to selection depends only on the additive effects of genes, h^2^ is the privileged estimated parameter (Visscher, Hill & Wray, 2008). In addition, quantitative genetic studies require keeping track of individual’s identity over the whole experiment either by rearing each individual separately or by marking them from birth to phenotypic measurement. This may be either very time and space consuming or technically challenging, in some species, and creates a practical limit to the obtainment of an adequate sample size. Therefore, for a given sample size, it seems crucial to optimize the crossing scheme (paternal or maternal half-sibs, number of families and offspring per family…) towards more statistical power, which depends on which components of the phenotypic variance are estimated (Lynch & Walsh, 1998).

Second, pedigree-free methods can release the constraints of keeping track of each phenotyped individual during the whole experiment. From a dataset of genotypes, one can either compute pairwise values of genetic relatedness or reconstruct the whole pedigree to incorporate in quantitative genetic models. These methods have been successfully used for quantitative genetic analyses in natural populations where pedigree information is generally not available except for long-term studies. In this field context, many simulation studies have explored their potential and limits, including quality and quantity of molecular markers and performance of relatedness coefficients (Visscher, Hill & Wray, 2008; Gay, Siol & Ronfort, 2013). Overall, performance of these methods rely mainly on the number and quality of molecular markers (Wang, 2006) and on relatedness composition of the sampled population (Csilléry, Johnson, Beraldi, *et al.*, 2006; DiBattista, Feldheim, Garant, *et al.*, 2009). Laboratory populations are closed systems where the relatedness composition can be optimized either by a total control of mating or with free mating of a chosen set of breeders. This latter option is particularly useful to maximize the probability of obtaining successful crosses when mating among designated individuals is not guaranteed, for example when mate choice is strong.

Third, the inheritance of various traits may be very complex. Since heritability estimates are based on the phenotypic resemblance of related individuals, they can be artificially inflated by resemblance caused by maternal effects (Kruuk & Hadfield, 2007). Using animal models, which are linear mixed models with the relatedness matrix as random factor, Wilson et al. (2005) estimated that maternal effects accounted for 21% of the total phenotypic variation in the birth weight of Soay sheep, compared to 12% for the heritability itself. Maternal effects can further be distinguished between environmental effects experienced by the mother, genetic variation among mothers and finally genotype-by-environment interactions. Accordingly, in Soay sheep, the maternal environmental effects and the maternal genetic effects represent respectively 11% and 12% of the phenotypic variation of birth weight (Wilson, Coltman, Pemberton, *et al.*, 2005)). To our knowledge, few studies have precisely quantified how heritability estimates can be biased by the presence of non-estimated maternal effects and even fewer have explored the precision of maternal effect estimates (but see Kruuk & Hadfield, 2007; Holand & Steinsland, 2016; De Villemereuil, Gimenez & Doligez, 2013). Even if the main motivation when considering maternal effect is to control this potential statistical nuisance in heritability estimates, maternal effects are also of considerable evolutionary interest to understand the evolution of traits. For example, theoretical models showed that maternal genetic effects represent an additional source of genetic variation which can affect the rate of trait evolution (Kirkpatrick & Lande, 1989).

In view of these considerations, and despite a vibrant field, some important methodological challenges still remain to be solved prior to address the quantitative genetics of a new model species: Can I omit maternal effects? What is the best statistical model to estimate the genetic basis of phenotypic traits? What are the sample size and structure of data required? Can pedigree-free approaches alleviate some of the technical constraints in quantitative genetics designs? We here addressed these four questions in the case study of the desert locust.

To this aim, we ran quantitative genetic analyses on an experimental dataset of body size, shape and color measured in late stages of a laboratory nature-derived population of the desert locust, *Schistocerca gregaria*, reared under controlled isolation conditions. We also used computer simulations to assess, along varying levels of heritability and maternal effects, the performance of two statistical animal models under various crossing schemes and relatedness inferences, derived from the experimental design. We finally interpreted phase trait data in the desert locust, illustrated how cautious one should be when interpreting this kind of data, and suggested directions for future research investigations.

Locusts are renowned for their nymphal marching bands and winged adult swarms that threaten food security in large areas (Sword, Lecoq & Simpson, 2010). This striking gregarious behavior is one aspect of the locust phase polyphenism, an extreme case of phenotypic plasticity. At low population densities, individuals tend towards a solitarious phenotype. On the contrary, at critically high population densities, locusts become gregarious. This fascinating phenotypic plasticity involves many traits (often called “phase traits”), amongst which behavior, morphometry, coloration, physiology and life-history traits (Pener & Simpson, 2009). The substantial variation in phase traits observed between natural populations, reared under standardized laboratory conditions might indicate that these traits have an evolutionary potential (Nolte, 1966; Chapuis, Estoup, Augé-Sabatier, *et al.*, 2008; Yerushalmi, Tauber & Pener, 2001; Botha, 1967; Schmidt & Albütz, 1996). However, the genetic contribution to phenotypic variance of key phase traits has never been assessed in locusts; and their potential to respond to selection is unknown. In this attempt, it would be informative to carry quantitative genetics experiments on both isolated and crowd-reared locusts, as phase polyphenism is a response to density. However, marking locusts throughout their development and successive molts is not feasible (Gangwere, Chavin & Evans, 1964), which makes methods based on molecular markers (*i.e.* pedigree-free methods) a promising alternative to estimate variance components of phase traits in crowd-reared locusts. Over and above that, for more than 50 years it has been known that parental rearing density also affect phase traits such as coloration and morphometry of hatchlings. Crowded parents tend to produce black and larger-headed hatchlings (and inversely for isolated parents), irrespective of the population density experienced by offspring during their development (see Table 1). Therefore, estimating maternal effects is of high relevance to the understanding of evolution of phase polyphenism.

**Table 1:**
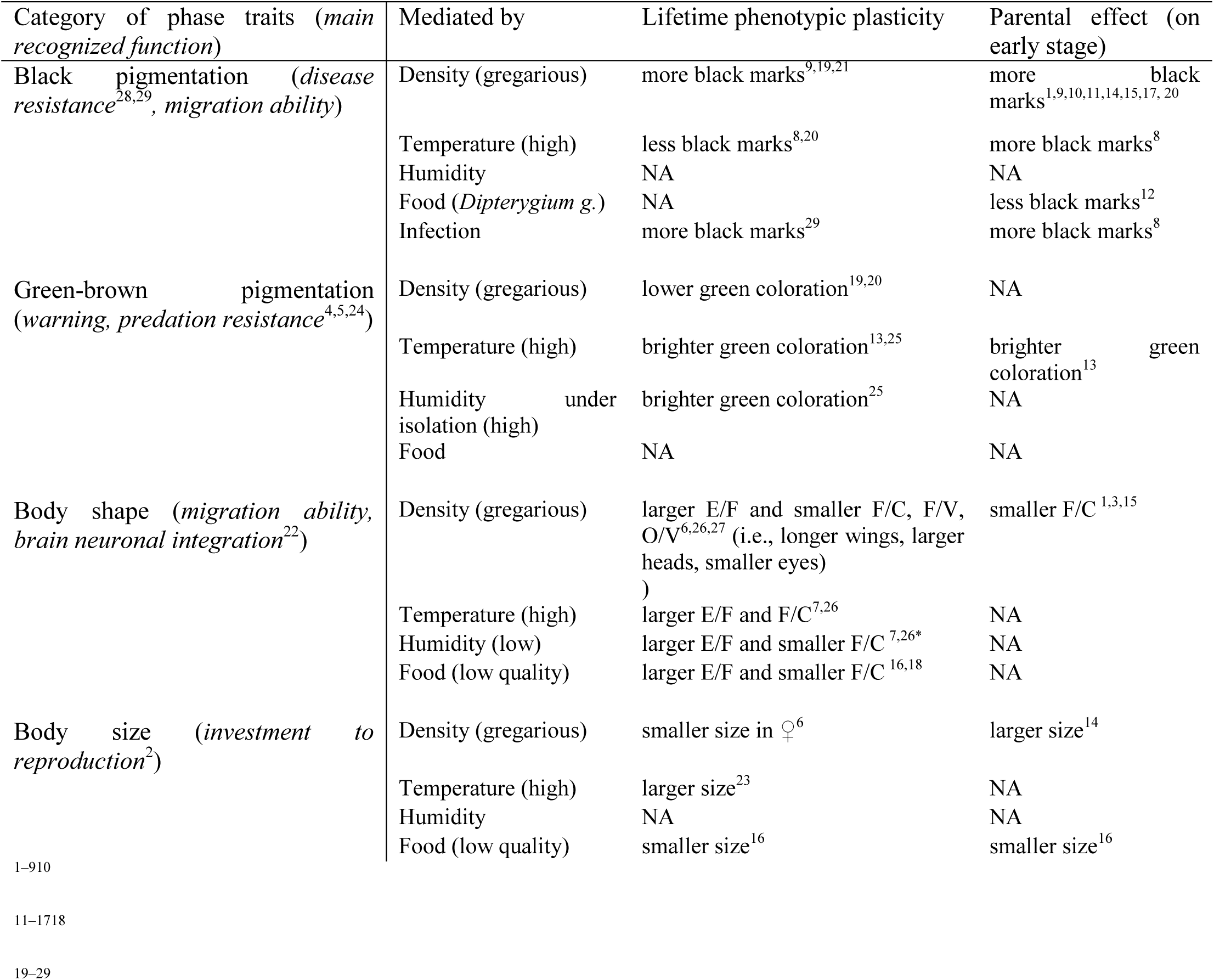
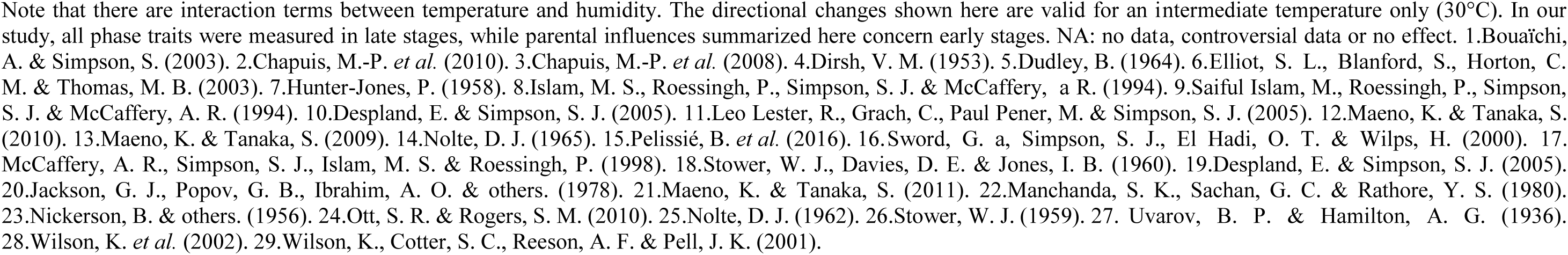
Literature-based evidence for environmental lifetime and parental effects on the phase traits measured in this study.

## Material and methods

### Quantitative genetics animal models

All quantitative genetics analyses were based on half-sib full-sib designs. We used two different kinds of animal models: Model 1 in which maternal effects are not specified (*i.e.* a naive model) and Model 2 which includes maternal effects (*i.e.* an informed model similar to equation 2 in De Villemereuil, Gimenez & Doligez, 2013).

**Model 1** only specifies a genetic effect as a random pedigree effect:

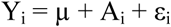

where Y_i_ is the phenotype of individual i, μ is the population mean, A_i_ is the individual’s additive genetic value, and ε_i_ is the random residual value. Hence, the total phenotypic variance (V_P_) was portioned into a variance attributed to additive genetic effects (V_A_) and a residual variance (V_R_) such that V_P_ = V_A_ +V_R_

**Model 2** specifies, in addition to pedigree, a random mother effect M_ki_ (environment of the mother k on individual i):

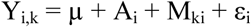

In this case, V_P_ = V_A_ + V_M_ + V_R_. From Model 1 and Model 2, we computed the narrow-sense heritability h^2^ = V_A_ / V_P_ and similarly, the maternal effect m^2^ = V_M_ / V_P_. This maternal effect includes maternal environmental effects, maternal genetic effects as well the interactions between genes and the environment (McAdam, Garant & Wilson, 2014). The estimation of the maternal genetic component would have required that individual mothers have female relatives in the dataset which is not the case in the studied half-sib/full-sib designs.

The random pedigree effect was computed from either a pedigree (pedigree-based method) or the relatedness between pairs of genotyped individuals (pedigree-free method). Additive genetic and maternal estimates were obtained by running univariate animal models using **Asreml-R** (Butler, Cullis, Gilmour, *et al.*, 2007). Standard errors for h^2^ and m^2^ were obtained by the delta method (Lynch & Walsh, 1998). P-values for the maternal effect were obtained by likelihood ratio tests (LRTs) between Model 1 and Model 2 whereas p-values for the pedigree effect were obtained by LRTs between Model 1 or Model 2 and the same model without the random pedigree effect (Wilson, Réale, Clements, *et al.*, 2010).

### Empirical data

#### Experimental design

Our laboratory population derived from fertilized desert locust females collected in the field (see Pelissie, Piou, Jourdan-Pineau, *et al.*, 2016 for further details). Locusts were maintained under isolated conditions for four subsequent generations. The fourth generation consisted in half-sib and full-sib families. The crossing scheme was 8 sires, mated to 2 to 3 females yielding to a total of 15 maternal families. The use of paternal half-sibs was dictated by our ambition to estimate maternal effects but also by the presence of multiple paternities in the desert locust (Seidelmann & Ferenz, 2002). For each maternal family, approximately 13 offspring were evenly distributed, right after hatching, between two temperature treatments: 28°C or 34°C. Temperature is known to affect phase traits (see Table 1) and may exert developmental constraints, susceptible to reveal genetic variation (Charmantier & Garant, 2005). A total of 486 hatchlings were selected and kept until adult molt. Larval mortality reduced the final sample size to 212 adult offspring. Known maternal effects were largely controlled with a homogenization of density, temperature and other main environmental drivers (e.g. humidity, food given *ad libitum*) (see Table 1 and Pelissie, Piou, Jourdan-Pineau, *et al.*, 2016 for further details on rearing isolation conditions).

#### Phenotypic measurements

We considered two commonly used sets of phase characteristics: fifth-instar larval coloration and adult morphometry (Pener & Simpson, 2009). Color differences between gregarious and solitarious desert locust larvae are the most noticeable phase change ((Nickerson & others, 1956; Pener & Simpson, 2009). Population density induces modification in the black patterning and in the green-brown coloration: solitarious late juveniles are typically green whereas gregarious late juveniles display a beige or brown background color with black pigmentation (Table 1 and references within). This is because all larvae have an integument of a beige or brown color, and either a black pigment, melanin, is deposited after ecdysis in the cuticle of the integument of gregarious insects, or a green pigment is produced from a yellow carotenoid and a blue bile pigment in the haemolymph of the integument of solitarious insects (Nolte, 1965). Thus, we measured the level of brightness directly correlated negatively to the level of black pigmentation and the percentage of green color which is a direct estimate of the green-brown polyphenism of an individual (see section 1.1 in the Appendix for details on methods and illustrations).

In adult, five morphometric ratios were considered: (i) the ratio of the length of the fore wing on the length of the hind femur (E/F) and (ii) the ratio of the length of the hind femur on the maximum width of the head (F/C), widely used for characterizing phase state in the field (Stower, Davies & Jones, 1960); (iii) the ratio of the length of the hind femur on the width of the vertex between eyes (F/V) and (iv) the ratio of the vertical diameter of eyes on the width of the vertex between eyes (O/V), considered as reliable indicators of phase change (Dirsh, 1953) (see section 1.2 in the Appendix for details on methods and illustrations). The values of these ratios changes toward gregarious adults with longer wings, larger heads and shorter eyes (Table 1 and references within).

In addition to larval coloration and adult morphometry, we considered two proxies of body size that varies with phase but in a sex-dependent manner. We measured the maximal larval weight (Pélissié et al. 2016) in the fifth-instar larvae and the length of the hind femur (F) in adults (with a low measurement error; e.g. Chapuis, Foucart, Plantamp, *et al.*, 2017). In adults of *S. gregaria*, solitarious females are larger than conspecific gregarious females, but solitarious males are slightly smaller than gregarious ones (Table 1 and references within). Therefore, the difference in body size between the females and the males is smaller in the gregarious than in the solitarious phase.

For each adult, we determined its sex to control for sexual dimorphism in body size and shape (Dirsh, 1953). We also recorded the number of larval molts, since between the third and fourth instars, desert locusts can undergo an extramolt that influences adult body size, E/F and F/C ratios (Pélissié, Piou, Jourdan-Pineau, *et al.*, 2016; Maeno, Gotoh & Tanaka, 2004). We summarized the larval color, adult body shape and size variables by extra-molting, sex, temperature in the section 1.3 in the Appendix. Details on maternal effects and functions of these density-mediated changes can be found in Table 1.

#### Quantitative genetics analyses

In order to remove non-genetic variation associated with known effects, we fitted sex, temperature, extramolting and their interactions as fixed effects in animal models (see section 1.4 in the Appendix). For each trait, we estimated the genetic component of phenotypic variance by running both Model 1 and Model 2. The random pedigree effect was estimated using the inverse of the additive genetic relationship matrix (A matrix) computed from a pedigree spanning 4 generations. Note that we obtained very similar results (data not shown) when using only the parental and offspring generations in the pedigree (*i.e.* 2 instead of 4 generations), indicating that its level of inbreeding was low (Lynch & Walsh, 1998). Finally, we also ran Model 1 replacing the pedigree by a marker-based relatedness matrix based on the genotyping of 96 offspring from the original dataset (all reared at 34°C) with a set of 16 microsatellite markers (SgM51, SgM92, SgM41, SgM74, SgM66, SgM96, SgM87, SgM88, SgM86, SgR36, DL09, SgR53, DL13, diEST-11, diEST-8 and diEST-40, Yassin et al. 2006, Kaatz et al. 2007, Blondin et al. 2013). Those last results were compared with analyses run on the same individuals but using the known pedigree instead of the pairwise relatedness values. To allow a complete comparison, we also ran Model 1 on the subset of individuals reared either at 28°C or at 34°C. However, the interaction between genotype and environment will not be treated further in this study.

### Simulated data

#### Simulation algorithm

The simulated phenotypic values were computed using Model 2 (which includes a maternal effect) and following Morrissey *et al.*, (2007): μ, the mean phenotype in the population, was arbitrarily set to 0; A_i_, the breeding value of the individual i, was normally distributed assuming additive genetic variance V_A_; M_k_, the maternal effect was normally distributed assuming variance V_M_, and ε_i_, the residual variation, was normally distributed with variance V_R_. To compute the breeding values A_i_ according to the simulated pedigree and V_A_, we used the **rbv** function from the R package **MCMCglmm** (Hadfield, 2010), which applies a Mendelian random deviation for each offspring.

#### Simulation of phenotypes

In every investigated scenario, we allowed the level of the heritability h^2^ and of the maternal effect m^2^ to vary among 4 fixed values: 0 (absence), 0.1 (low level), 0.3 (moderate level) and 0.5 (high level). Those values are realistic in regard to previous studies in insects and, more generally, in other animals (Mousseau & Roff, 1987; Houle, 1992; Visscher, Hill & Wray, 2008 for h^2^ values; Räsänen & Kruuk, 2007; Wilson, Coltman, Pemberton, *et al.*, 2005 for m^2^ values). They were obtained by setting the total phenotypic variance V_P_ to a fixed value while allowing V_A_, V_M_ and V_R_ to vary. We generated every possible combination of h^2^ and m^2^, thus leading to the comparison of 16 different phenotypic scenarios.

#### Simulation based on our experimental design

For each combination of h^2^ and m^2^, we simulated 1,000 phenotypic datasets based on our experimental design, *i.e.* with exactly the same pedigree and the same subset of phenotyped individuals (Morrissey, Wilson, Pemberton, *et al.*, 2007).

#### Simulation based on refined crossing schemes

We tested the sensitivity of estimation to various paternal half-sib/full-sib designs (see section 2.1 in the Appendix for parameter values of each test crossing scheme). We first simulated a design very close to our actual experimental design: it resulted in a crossing scheme of 8 sires, 2 dams by sires and 13 offspring per dams, for a total of 208 offspring (CS3). We then used this half-sib/full-sib design as a reference to derive 13 more crossing schemes with varying numbers of sires (S), dams by sire (D), and of offspring (O) per family (F = D × S), for a total sample size (N) of 208 offspring. This set of crossing schemes allowed us to compare the distinct effects of family size and the crossing scheme on our ability to accurately detect h^2^ and m^2^. Finally, we tested for the effect of doubling the total sampling size by simulating a 15^th^ dataset involving 416 offspring (CS15). This crossing scheme was derived from one of the best performing crossing scheme (see below) and consisted of 8 sires, 13 dams by sire and 4 offspring per dam instead of 2 (CS12). For each crossing scheme and for each combination of h^2^ and m^2^, we simulated 400 phenotypic datasets.

#### Pedigree-free approaches

First, we ran the same type of simulations but replacing pedigree-based relatedness by marker-based relatedness, on 2 crossing schemes: our experimental design (as if the 212 individuals had been genotyped) and the crossing scheme CS15 with 416 individuals. In each design, we tested 2 realistic sets of microsatellite markers: 16 (the set used for genotyping in our experimental design), or 29, that is the maximum number of markers available for the desert locust (Kaatz, Ferenz, Langer, *et al.*, 2007; Yassin, Heist & Ibrahim, 2006; Blondin, Badisco, Pagès, *et al.*, 2013). Pairwise relatedness based on microsatellite markers were computed using the coefficient introduced by Loiselle et al. (1995). We used the R package **pedantics** to simulate molecular genotypes based on a selection of markers and a given pedigree (Morrissey & Wilson, 2010) and the R package **Ecogenetics** to compute Loiselle relatedness coefficients based on the desert locust microsatellite markers (Roser, Vilardi, Saidman, *et al.*, 2015). The analyses were processed with Model 1 only since maternal identity could not be inferred from molecular relatedness. We explored 4 scenarios with respective h^2^ values of 0, 0.1, 0.3 and 0.5 and simulated 400 relatedness matrices (based on Loiselle coefficients) and phenotypic datasets per scenario.

#### Performances of simulated datasets

We evaluated the performance of the animal models, crossing schemes and pedigree-free methods using four criteria applied to all simulations within one scenario: i) the mean values of h^2^ and m^2^ estimates, ii) the 95% confidence intervals; which inform on bias and dispersion, respectively, iii) the average of the root mean square error (RMSE) between simulated and estimated values (Bolker, 2008) and iv) the power to detect either pedigree or maternal effect computed as the percentage of simulated datasets that gave a significant pedigree effect or maternal effect (when included). To compare the simulated crossing schemes, we tested the influence of dam-to-sire ratio (D:S), of the number of offspring per family (O) and of their interaction, on the RMSE (of h^2^ and m^2^) and on the statistical powers to detect pedigree and maternal effect), using linear models with the levels of h^2^ and m^2^ (respectively) as covariate.

## Results

### Empirical dataset on phase traits of the desert locust

Heritability and maternal effects estimates computed from the whole desert locust dataset are given in Table 2. In the naïve model (Model 1), body size traits, pronotum coloration traits, and the ratio of the femur length over the head width had significant h^2^ estimates (0.71 ≥ h^2^ ≥ 0.18). Interestingly, the same five traits were still significantly heritable when considering insects reared at the low temperature of 28°C. Conversely, none of the heritabilities turned significant at the high temperature of 34°C (see section 1.5 in the Appendix).

**Table 2:** Estimated genetic parameters for morphological and colour traits of the desert locust estimated from a model including either pedigree only (model 1) or pedigree and mother (model 2) as random effects. We used the whole experimental dataset and the real pedigree. We presented values for phenotypic mean and variance (computed on raw data), additive genetic variance (V_A_), variance associated with maternal identity (V_M_) and residual variance (V_R_), heritability (h^2^), maternal effect (m^2^) and their standard errors (SE), p-values of the pedigree effect and maternal effect. *Brightness*: Level of brightness, which is inversely related to the level of black pigmentation; *%Green*: Percentage of green color; *E*: Length of the fore wing; *F*: Length of the hind femur; *C*: Maximum width of the head; *H*: Height of the pronotum; *P*: Length of the pronotum; *O*: Vertical diameter of eyes; V: the width of the vertex between eyes.

| Trait | Mean | Variance | Model 1 with pedigree |  |  |  |  | Model 2 with pedigree and mother |  |  |  |  |  |  |  |  |
| --- | --- | --- | --- | --- | --- | --- | --- | --- | --- | --- | --- | --- | --- | --- | --- | --- |
|  |  |  | V<sub>A</sub> | V<sub>R</sub> | h<sup>2</sup> | SE(h<sup>2</sup>) | p-value (pedigree) | V<sub>A</sub> | V<sub>M</sub> | V<sub>R</sub> | h<sup>2</sup> | SE(h<sup>2</sup>) | p-value (pedigree) | m<sup>2</sup> | SE (m<sup>2</sup>) | p-value (mother) |
| %Green | 0.442 | 1.29E-03 | 1.03E-03 | 4.20E-04 | 0.71 | 0.28 | 0.00 | 1.45E-09 | 3.32E-04 | 9.06E-04 | 0.00 | 0.00 | 1.00 | 0.27 | 0.12 | 0.11 |
| 1/Brightness | 0.008 | 1.85E-06 | 6.74E-07 | 1.12E-06 | 0.38 | 0.21 | 0.00 | 2.28E-12 | 2.88E-07 | 1.42E-06 | 0.00 | 0.00 | 1.00 | 0.17 | 0.09 | 0.24 |
| E/F | 2.041 | 5.41E-03 | 2.00E-04 | 4.62E-03 | 0.04 | 0.08 | 0.53 | 2.00E-04 | 2.54E-09 | 4.62E-03 | 0.04 | 0.08 | 0.70 | 0.00 | 0.00 | 1.00 |
| F/C | 3.869 | 4.55E-02 | 8.07E-03 | 3.56E-02 | 0.18 | 0.14 | 0.04 | 6.26E-08 | 2.90E-03 | 3.91E-02 | 0.00 | 0.00 | 1.00 | 0.07 | 0.05 | 0.13 |
| F/V | 14.104 | 3.18E+00 | 1.90E-01 | 2.65E+00 | 0.07 | 0.09 | 0.15 | 1.46E-02 | 9.98E-02 | 2.72E+00 | 0.01 | 0.17 | 0.98 | 0.04 | 0.09 | 0.72 |
| O/V | 2.100 | 8.46E-02 | 2.70E-03 | 6.75E-02 | 0.04 | 0.06 | 0.39 | 2.70E-03 | 5.55E-09 | 6.75E-02 | 0.04 | 0.06 | 0.80 | 0.00 | 0.00 | 1.00 |
| F | 3.393 | 1.34E-02 | 2.57E-03 | 9.84E-03 | 0.21 | 0.14 | 0.01 | 1.76E-08 | 1.03E-03 | 1.10E-02 | 0.00 | 0.00 | 1.00 | 0.09 | 0.06 | 0.30 |
| Max larval weight | 1.708 | 2.15E-01 | 1.39E-02 | 1.73E-02 | 0.45 | 0.20 | 0.00 | 4.12E-03 | 3.62E-03 | 2.20E-02 | 0.14 | 0.46 | 0.78 | 0.12 | 0.20 | 0.63 |

Adding a maternal effect in the statistical model (*i.e.* the informed model, Model 2) strongly lowered additive genetic variances for most traits, due to large maternal variances (V_m_) compared to additive genetic variances. Nevertheless, none of these maternal effects were found to be significant (p-values ≥ 0.11), due to large standard errors of m^2^ estimates (SE ≥ 0.05). Conversely, the morphometric ratios E/F and O/V showed almost-null V_m_ and null m^2^ values, leading to the same values as in Model 1 for V_A_, h^2^ and SE(h^2^), although associated p-values were largely increased in Model 2.

Using the pedigree-free method with 16 microsatellite markers and on a subset of individuals measured at 34°C, we obtained variance and heritability estimates in the same order of magnitude as when analyzing the same subset using the real pedigree for most traits. However, E/F has larger additive genetic variance with the pedigree-free method than when using the pedigree (Table 3). With both methods, brightness was found significantly heritable (Table 3).

**Table 3:**
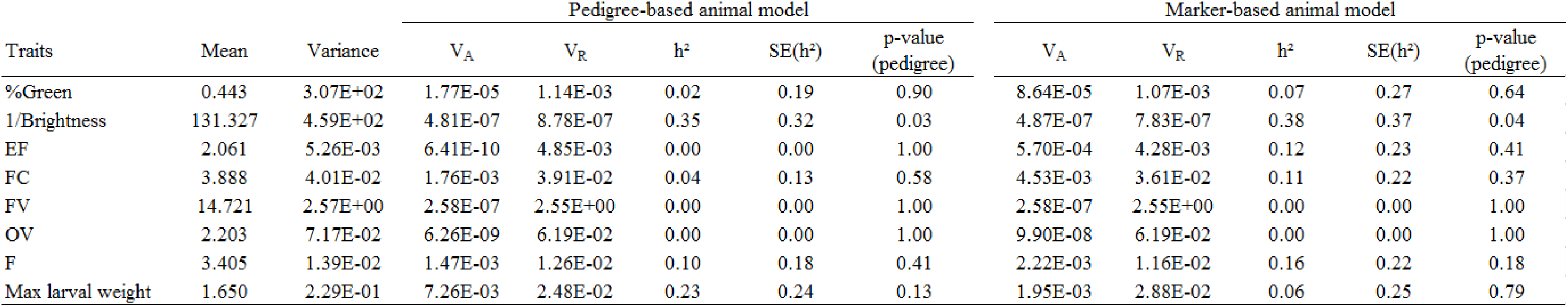
Estimated genetic parameters for morphological and colour traits, estimated from either the real pedigree or molecular relatedness computed from 16 microsatellite markers. We used a subset of the experimental dataset constituted of 57 larvae and 96 adults, all reared at 34°C, and a statistical model including pedigree only as random effect (Model 1 in text). The fixed effects were sex * extramolting for all traits. We presented values for phenotypic mean and variance (computed on raw data), additive variance (V_A_), residual variance (V_R_), heritability (h^2^), its standard error (SE(h^2^)), and p-value associated with the pedigree effect. *Brightness*: Level of brightness, which is inversely related to the level of black pigmentation; *%Green*: Percentage of green color; *E*: Length of the fore wing; *F*: Length of the hind femur; *C*: Maximum width of the head; *H*: Height of the pronotum; *P*: Length of the pronotum; *O*: Vertical diameter of eyes; V: the width of the vertex between eyes.

### Simulation based on our experimental design

In the absence of simulated maternal effects (m^2^ = 0; Fig. 1 first column), Model 1 performed better than Model 2 in estimating heritability. First, h^2^ estimates were biased downward only in Model 2 (e.g. a simulated h2 of 0.3 was estimated in average at 0.20). Second, both statistical models led to large dispersion in h^2^ estimates that increased with simulated h^2^ values, and RMSE values were close to h^2^ values (e.g. 0.16 for a simulated h^2^ of 0.3 in Model 1). Finally, in Model 1, the power to detect a true pedigree effect was low for low simulated h^2^ values (*i.e.* 30.5% for h^2^ = 0.1), satisfying for intermediate simulated h^2^ values (*i.e*. 82% for h^2^ = 0.3) and very high for the highest simulated h^2^ values (*i.e.* 95.9% for h^2^ = 0.5). Conversely, in Model 2, the statistical power stayed very low even when simulated heritability was the highest (*i.e.* 11.2% for h^2^ = 0.5).

**Figure 1:**
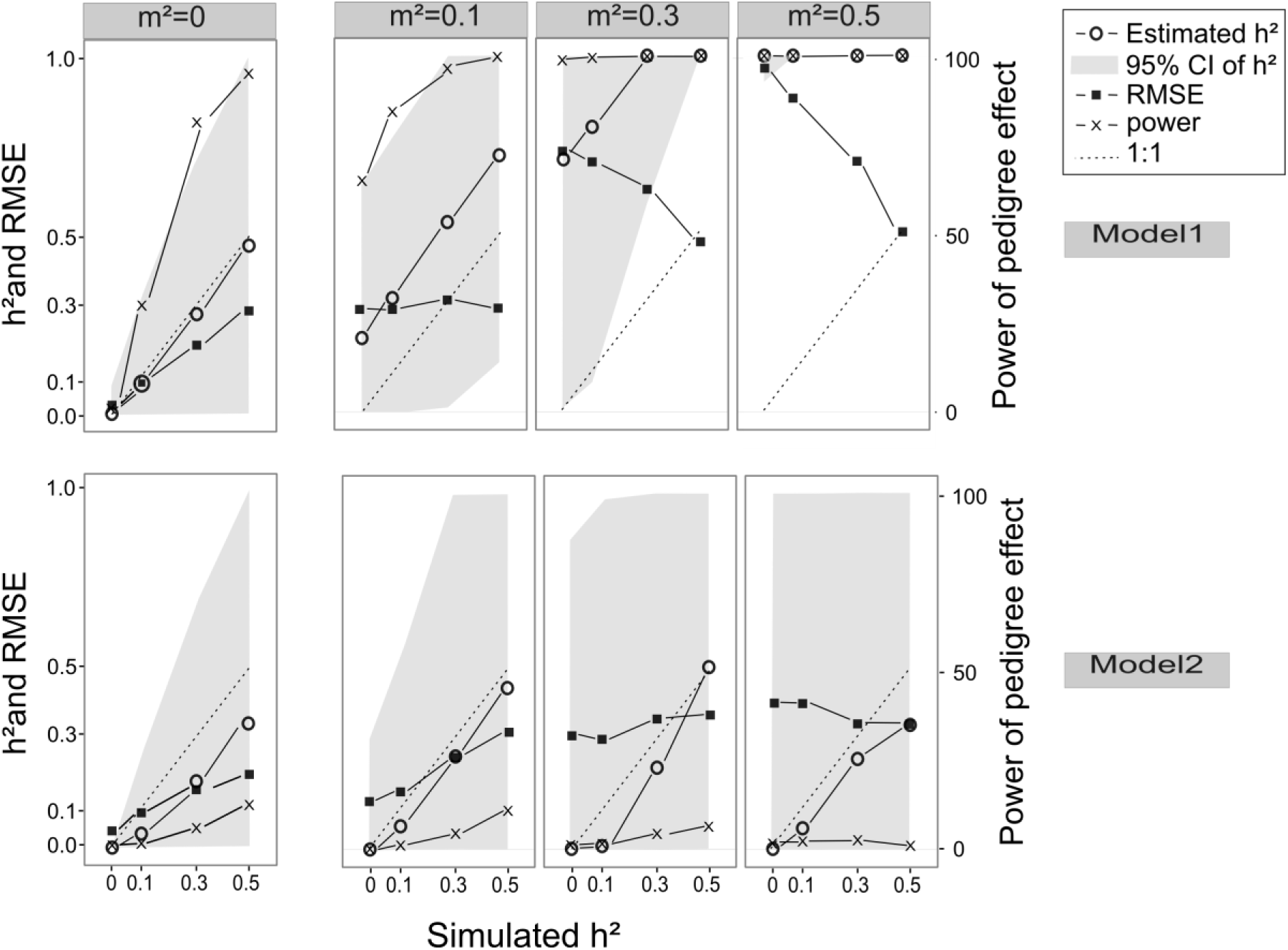
Performance of heritability estimates evaluated from simulation datasets based on our experimental design. We show mean estimate (h^2^) and 95% confidence interval (empty circles and grey area, respectively), root mean square error (RMSE) (black squares) and percentage of simulations with significant pedigree effect (crosses) (y-axis) as a function of simulated h^2^ (x-axis) and maternal effects (horizontal panels). We used either Model 1 (without specified maternal effect, top panels) or Model 2 (specifying a maternal effect, bottom panels).

In the presence of simulated maternal effects, the h^2^ estimates became highly biased upward with Model 1, reaching values of 1 for m^2^ = 0.5 whatever the simulated h^2^, or for m^2^ = 0.3 when simulated h^2^ was high (≥ 0.3) (Fig. 1, right upper panels). Accordingly, Model 1 generated significant pedigree effects in all simulations for maternal effects ≥ 0.3, even when the simulated heritability was null (*i.e.* 100% of false positives). Adding a simulated maternal effect in Model 2 induced a downward bias of the same magnitude but a greater dispersion of the h^2^ estimates, with even 95% CI covering the whole space when maternal effects were large (*i.e.* ≥ 0.3) (Fig. 1, right lower panels). The RMSE values for h^2^ estimates were however lower with Model 2 than with Model 1. In Model 2, the power for detecting a pedigree effect of any level was always very low (< 5%).

Estimation of a maternal effect with Model 2 showed a downward bias that decreased with higher simulated h^2^ (e.g. a simulated m^2^ of 0.3 was estimated in average in the range of 0.2-0.29; Fig. 2). Whatever the simulated h^2^ values, there was a large dispersion in estimates increasing with simulated m^2^ values. As for heritability estimates, RMSE values for maternal effects increased with the simulated m^2^ (0.17 to 0.20 for a simulated m^2^ of 0.3). The power to detect a maternal effect was low and just reached 50% when maternal effect was 0.5.

**Figure 2:**
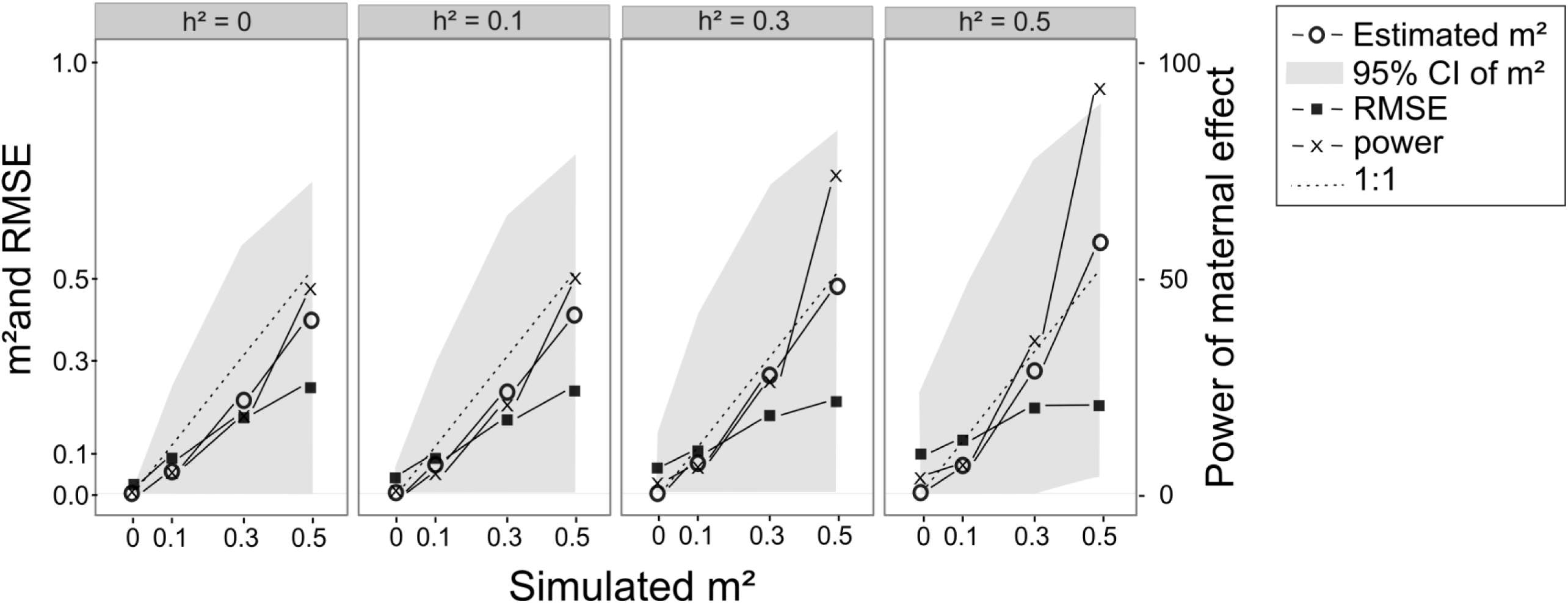
Performance of maternal effects estimation evaluated from simulation datasets based on our experimental design. We show mean estimates (m^2^) and 95% confidence intervals (empty circles and grey area, respectively), root mean square error (RMSE) (black squares) and percentage of simulations with significant maternal effect (crosses) as a function of simulated m^2^ (x-axis) and simulated h^2^ (panels). Estimates were obtained with Model 2.

### Simulation datasets on the varying crossing schemes

In the absence of maternal effects, the use of Model 1 on crossing schemes with more sires than dams by sires (D:S < 1) yielded slightly smaller RMSEs for h^2^ estimates ((F_1,55_=9.52, p-value=0.003) but did not improve the statistical power to detect a pedigree effect, (Fig. 3a). Conversely, a higher number of offspring per females (*i.e.* fewer families) did not impact RMSE values but yielded greater power for detecting a pedigree effect (F_1,55_=9.54, p-value=0.003, Fig. 3b). In presence of maternal effects, refining crossing schemes by the number ratios of dams on sires or of offspring on families did not help sorting out the upward bias and over power in heritability estimation of Model 1.

**Figure 3:**
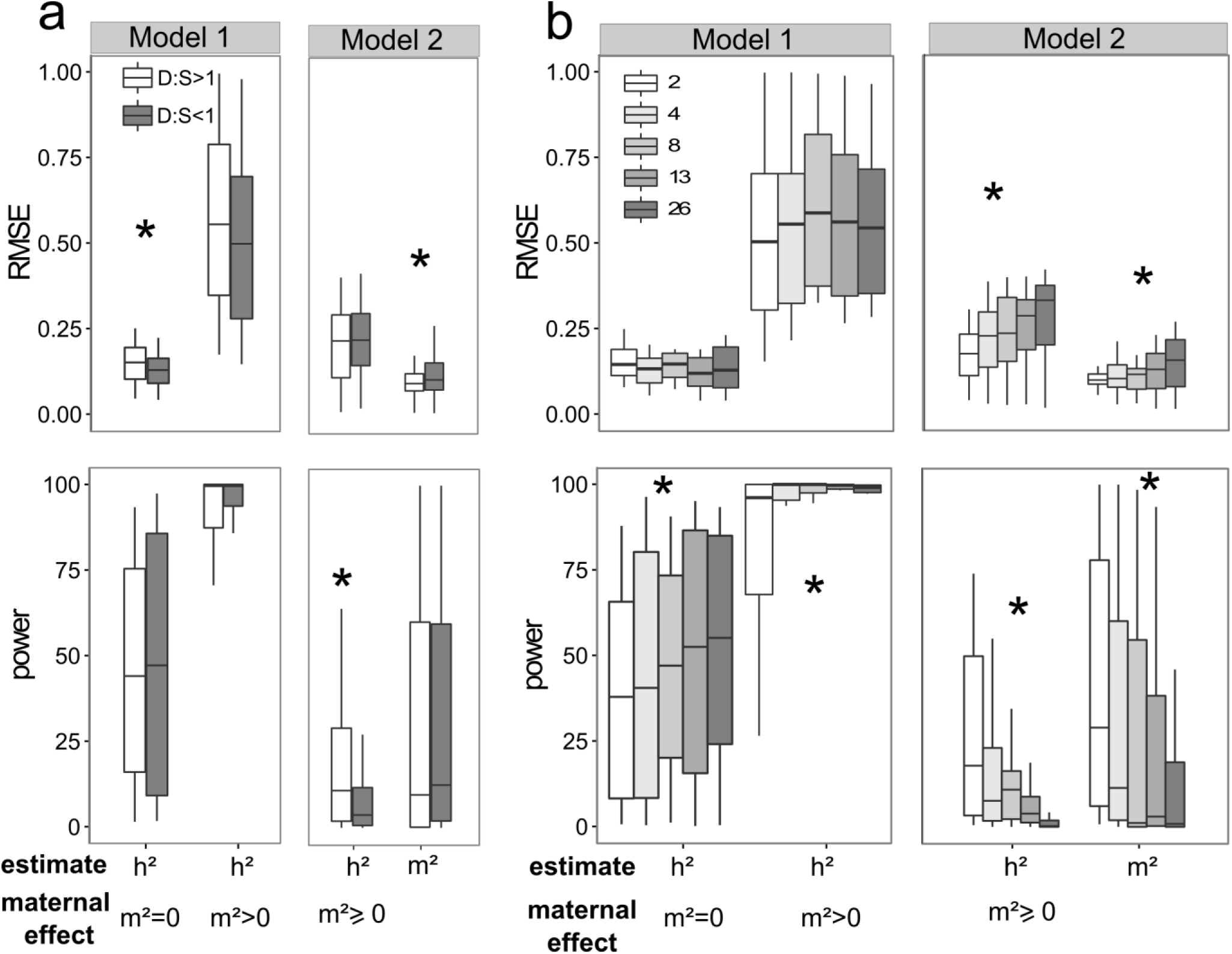
Effects on performance of h^2^ and m^2^ estimation of (a) the relative number of sires and dams by sire (larger: D:S<1 or smaller: D:S>1) and (b) the number of offspring per family for a given total sample size. We plotted the mean RMSE (top) and the mean power to detect h^2^ and m^2^ effects (bottom), both calculated over all values of h^2^ or m^2^ in relation with animal model (indicated above each panel) and absence or presence of maternal effect (indicated at the bottom of the graphs). 2, 4, 8, 13 and 26 are the numbers of offspring per family. * denotes a significant effect within a block.

Using Model 2 (with maternal effects ≥0), the power for detecting a pedigree effect was significantly greater in designs with D:S > 1 (F_1,223_= 11.56, p-value<10^−3^, Fig. 3a). RMSE for maternal effect estimates (m^2^) were significantly lowered with such crossing schemes (F_1,223_=6.89, p-value=0.001), but the power to detect them remained unaffected. Finally, increasing the number of families instead of offspring per female significantly increased both the power to detect heritability and maternal effect (F_1,239_= 100.17 p-value<10^−3^ and F1=105.15 p-value<10^−3^, respectively) while decreasing RMSE values (F_1,239_=52.33 p-value<10^−3^, and F_1,239_=57.67 p-value<10^−3^ respectively, Fig. 3b). Accordingly, one of the crossing schemes with the highest global performance in an informed model was composed of 8 families with 13 dams per sire and 2 offspring (CS12). This crossing scheme did not improve the small downward bias on h^2^ estimation but markedly decreased the variance in h^2^ estimation (i.e. 95% CI and RMSE criteria; see section 2.2 in the Appendix for details). This resulted in an increased power to detect a pedigree effect that could reach 62-74% for large maternal effects (i.e. m^2^=0.5) whereas it reached a limit of 11% under the crossing scheme mimicking our experimental design (CS3; Figure 1). As for maternal effects, this crossing scheme of a relative high numbers of families and of dams per sire allowed an unbiased estimation with a lowered variance (RMSE values ≤ 0.12 and narrower 95% CI) and an increased statistical power reaching 100% in the best case (m^2^=0.5).

Finally, we explored the gain in performance for a sample of a larger size. To this aim, we selected the crossing scheme CS12 with a high global performance, doubled the number of offspring within families (*N* = 416) and ran additional simulations with this new crossing scheme (CS15). In the absence of a maternal effect, Model 1 showed good performances, with a slight increase in power to detect a pedigree effect and a slight decrease in RMSE values (Fig. 4, left upper panel). The performance of Model 2 in h^2^ estimation was increased, with a reduced downward bias and augmented power in h^2^ estimates, but still lower than in Model 1 (Fig. 4, left bottom panel). In the presence of maternal effects, the performance of Model 1 to estimate the pedigree effect was still poor in line with previous simulations, without improvements of the strong overestimation and high number of false positives (Fig. 4, upper right panels). With Model 2, h^2^ estimation was not biased downward anymore with this large sample size design and the power to detect a pedigree effect considerably increased, though still low (≤ 72%) when maternal effects were high (≥ 0.3; Fig. 4, lower right panels). In comparison with the same type of crossing scheme with twice lower sample size, maternal effects were estimated more precisely (narrower 95% CI) and with greater power (Fig. 5).

**Figure 4:**
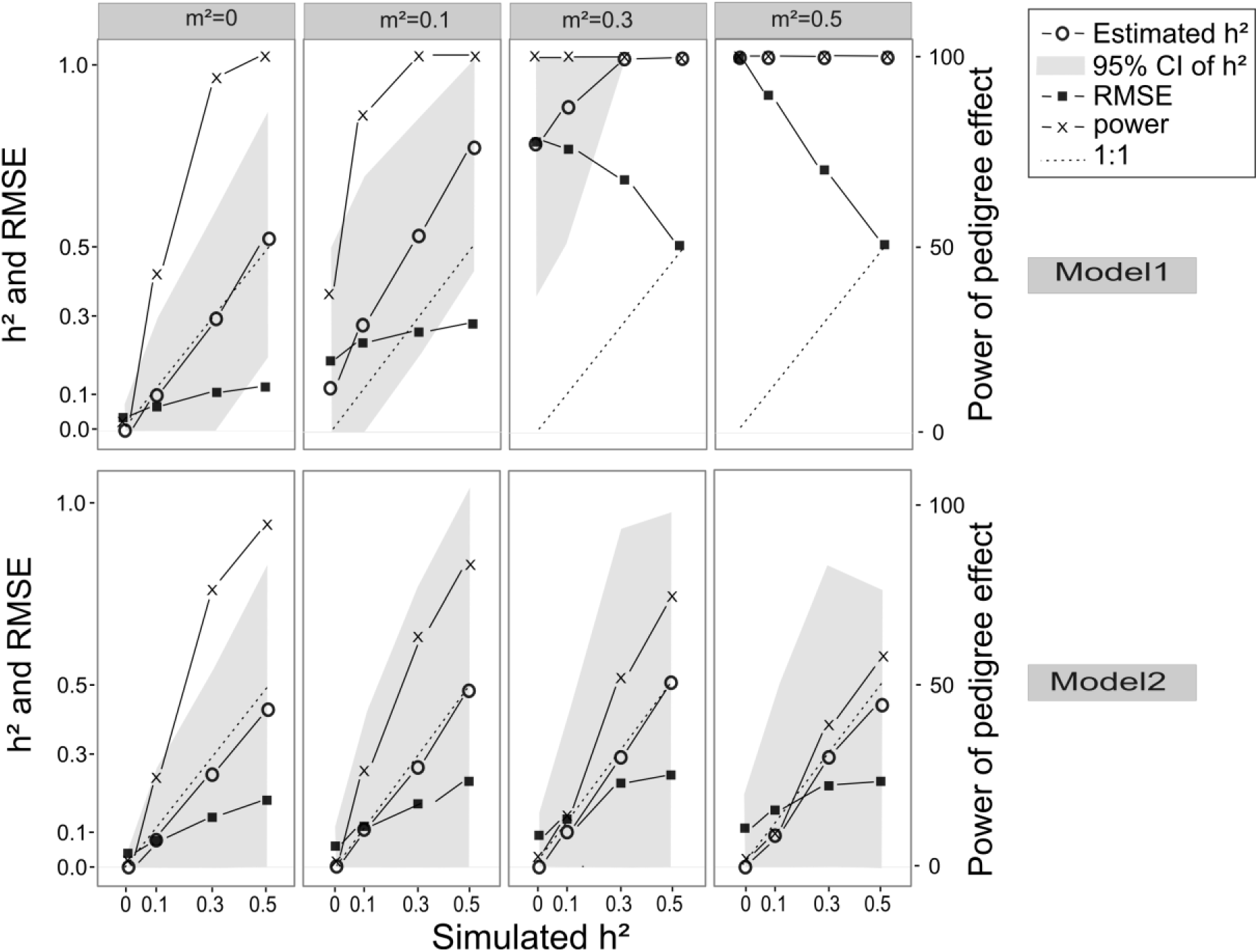
Performance of heritability estimation evaluated from simulation datasets on the best crossing scheme (CS15, 416 measured offspring). We show mean estimate (h^2^) and 95% confidence interval (empty circles and grey area, respectively), root mean square error (RMSE) (black squares) and percentage of simulations with significant pedigree effect (crosses) (y-axis) as a function of simulated h^2^ (x-axis) and maternal effects (horizontal panels) We used either Model 1 (without specified maternal effect, top panels) or Model 2 (specifying a maternal effect, bottom panels).

**Figure 5:**
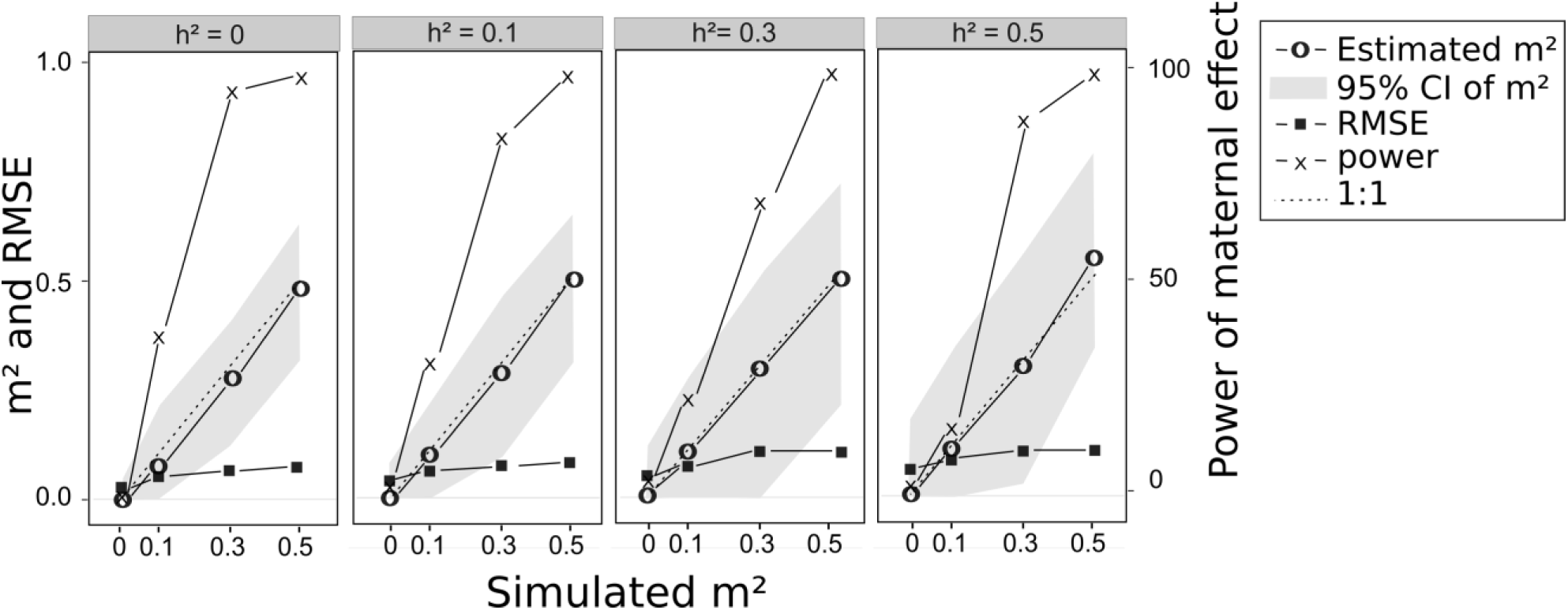
Performance of maternal effects estimation evaluated from simulation datasets on the best crossing scheme (CS15, 416 measured offspring). We show mean estimates (m^2^) and 95% confidence intervals (empty circles and grey area, respectively), root mean square error (RMSE) (black squares) and percentage of simulations with significant maternal effect (crosses) as a function of simulated m^2^ (x-axis) and simulated h^2^ (panels). Estimates were obtained with Model 2.

### Simulation datasets with pedigree-free method

Overall, simulations based on our experimental dataset and on the large crossing scheme (CS15) showed very similar outcomes (Fig. 6). Using relatedness values computed from genotypes of 16 or 29 microsatellite markers yielded very close performances of h^2^ estimation, both in terms of RMSE and power to detect a pedigree effect. Pedigree-free methods performed reasonably well when compared to using the full pedigree, showing only a slight 2-10% decrease in power, and a 30% increase in RMSE in the worst case, i.e. the smallest number of microsatellite markers and a high simulated heritability (h^2^ = 0.5). This increase in RMSE was explained by a downward bias in h^2^ estimates when using microsatellite markers compared to using the full pedigree (results not shown).

**Figure 6:**
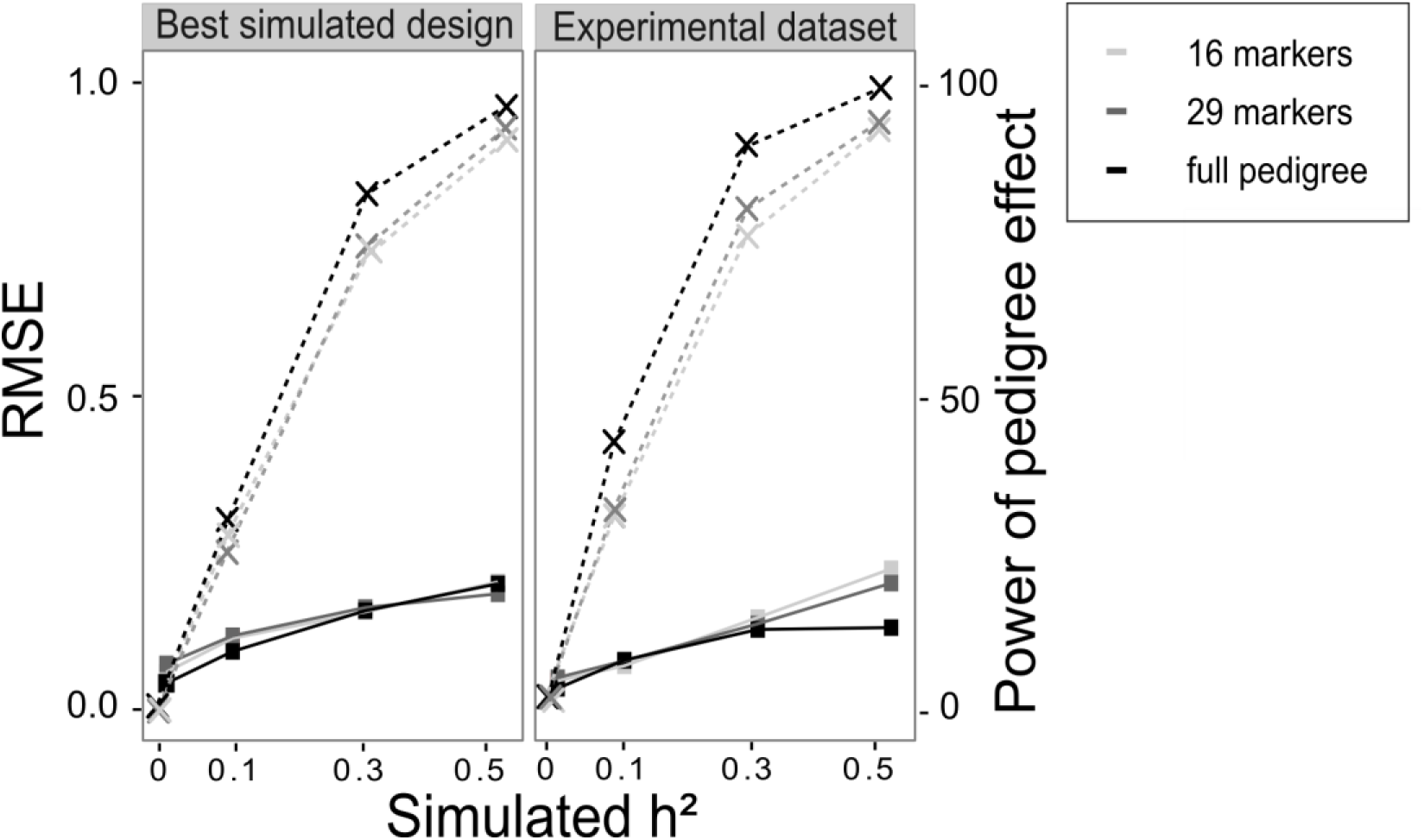
Performance of heritability estimation evaluated from simulation datasets on marker-based pedigree free method. We analyzed two different numbers of microsatellite markers (16 or 29), and the pedigree method is also shown as a reference. Simulations were performed on two designs: the best simulated design (CS15, 416 measured offspring) and our experimental design. Performance was evaluated by the root mean square error (RMSE) (squares and solid lines) and the power to detect a pedigree effect (crosses and dashed lines).

## Discussion

Statistical limitations in quantitative genetics studies may compromise to draw firm conclusions about the genetic basis of the traits under study. The present study used computer simulations to examine the validity and limits of a standard quantitative genetics experiment, in the context of the density-dependent phase polyphenism, partly transmitted by maternal effects. We looked at the performance of animal models in disentangling heritability and maternal effects, and how these performances were affected by the crossing scheme and the relatedness inference. We interpreted phase trait data in the desert locust in the light of the simulation results and recommended methodological directions for future research.

### Performance of a naïve model (Model 1)

In absence of maternal effects, a naive model (without any specified maternal effect) outperformed an informed model (Model 2) in heritability estimation, whatever the type and sampling size of crossing schemes. Our experimental half-sib/full-sib design led to unbiased estimation with the naïve model as well as a satisfying power, except for low levels of heritability (e.g. h^2^=0.1) (Fig. 1). In such situation, crossing schemes with more sires than dams by sires showed the greatest performances to estimate heritability (Fig. 3). This result echoes the classical calculation of h^2^ in half-sib/full sib design analyses where h^2^ is directly derived from the sire variance; therefore more sires should give greater precision to h^2^ (Lynch & Walsh, 1998). This kind of crossing scheme might also be advantageous in species where it is easier to use a large number of males mated to few females each. This is the case for the desert locust, whose mating can last several hours to several days, strongly decreasing the potential number of female partners per males (Uvarov, 1966). In addition, in the naive model, the power to detect pedigree effect was greater with a larger number of offspring per female but this was not accompanied by any improvement in RMSE values (Fig. 3). In conclusion, a crossing scheme close to the one we used for the acquisition of experimental data on phase traits of the desert locust is relevant for the estimation of heritability in absence of a maternal effect (see the summary guideline in Table 4). A standard sample size should provide robust information on moderate and high heritability traits, even if larger effort would improve the power and precision of estimation.

However, in the naive model, the presence of a maternal effect strongly inflated heritability estimates (and statistical power), thus producing a large number of false positives, whatever the type and sample size of crossing schemes (Fig. 2). Two previous studies, using the same restricted maximum likelihood method, also warned about the overestimation of heritability estimates when maternal effects are not specified in the animal model (De Villemereuil, Gimenez & Doligez, 2013; Kruuk & Hadfield, 2007). In Kruuk and Hadfield (2007), the overestimation was large, as in our study, with a mean estimated h^2^ of 0.52 (bird system) or even 0.6 (ungulate system) for a simulated h^2^ of 0.3 and m^2^ of 0.2. In comparison, Villemereuil et al. (2013) found smaller bias in h^2^ caused by maternal effect: for instance, they obtained a simulated h^2^ of 0.2 for a simulated h^2^ of 0.1 and m^2^ of 0.45. This lower effect of maternal effect in the h^2^ estimates may be due low levels of m^2^ in their simulations (Villemereuil et al (2013)).

### Performance of an informed model (Model 2)

Since maternal effects lead to overestimate heritability in a naive model, under their suspicion, it seemed appropriate to consider an informed model (specifying maternal effects). With our experimental dataset, h^2^ estimates were shown to be little biased downward but, the power to detect a pedigree effect became null or very low (< 11%; Fig. 2). The low performance in h^2^ estimation was improved by an increased number of families (instead of a large number of offspring per female) and a number of dams by sire greater than a number of sires (Fig. 3). The former result is in agreement with theoretical formulae of sampling error and power of heritability estimates (Lynch & Walsh, 1998) whereas the latter is probably linked to the greater precision of estimation of the maternal effect with larger numbers of females per male. Villemereuil et al. (2013) showed that parent-offspring regression, restricted maximum likelihood (tested here), and Bayesian methods (both using an informed model) performed similarly in estimating heritability in the presence of a maternal effect. However, parent-offspring regression requires measurements of both parents and offspring and Bayesian method gives even more biased results with small sample size (De Villemereuil, Gimenez & Doligez, 2013).

Our simulations also showed that the informed model estimated maternal effects more precisely than the heritabilities. However, optimized crossing schemes (Fig S4, Appendix) or large sample sizes (about 400 offspring, Fig 5) are needed to detect maternal effects with sufficient power. Otherwise, LRT between nested models should be used with caution to decide whether maternal effects are significant and which model to use. With the Bayesian approach, Holand and Steinsland (2016) demonstrated that using the Deviance Information Criterion (DIC, a generalization of the Akaike Information Criterion) to compare naive and informed models also required a substantial maternal effect (equal to half the heritability), even with a very large sample size (*N*=1025).

### Some recommendations regarding models and designs

When sample size or crossing scheme are practically constrained, our simulations confirmed that specifying the right animal model is crucial to have sufficient power and reliable estimates of pedigree and maternal effects: omitting a maternal effect in the statistical model generates overestimation of heritability and false positives whereas inappropriately specifying a maternal effect dramatically wipes out the power of analyses. Since maternal effect estimation is more accurate than pedigree effect estimation, we advise to first inform on the maternal effect using an informed model, and then decide, from the obtained *P*-values and estimate values, which model should be used. Note that comparing outputs of both statistical models may also provide an indication on the absence of a maternal effect, since in such a case, both models should give congruent h^2^ estimates. However, in the case where a maternal effect is estimated to be present, interpreting results must be done with caution since the power to detect a pedigree effect would remain low and the study might be inconclusive (see the summary guideline in Table 4). Furthermore, in the case where a maternal effect is estimated to be absent, the use of a naïve model might be done with a sub-optimal crossing scheme, as requirements of this model are opposite in relative numbers of dam by sires to sires and of offspring per family to family. Thus, without prior knowledge on the presence of a maternal effect, the best option to estimate h^2^ might be to favor the greatest number of families and a balanced number of sires and dams by sires.

### Use of pedigree-free methods

Analyzing big datasets with strong relatedness structure, in order to get good detection power and accurate estimates of h^2^ and m^2^, implies being able to rear a lot of individuals in private boxes (to identify them if they cannot be marked) and to manipulate a lot of mating pairs. Private boxes represent an obvious constraint on experimental designs: more individuals mean more effort in sampling, rearing and manipulations. In addition, creating lots of mating pairs can prove to be challenging, especially in species where successful mating is not straightforward, for example if sexual selection is strong. In addition, in the context of phase polyphenism, manipulating rearing density of locust would be a requirement to carry comprehensive quantitative genetics experiments. Rearing individuals in group cages would alleviate these limitations, both reducing constraints on mating (increasing the number of families and the half-sib/full-sib structure) and allowing studying more individuals effortlessly. However, this removes the possibility to use a classical pedigree since mating pairs cannot be known exhaustively, calling for the use of pedigree-free methods.

Our results showed that, using a matrix of molecular pairwise relatedness computed at16 microsatellite markers might be sufficient to obtain reliable heritability estimates, despite slight decrease and downward bias in estimation precision in comparison with the use of a full pedigree. These results are more encouraging than those from most simulation studies in a context of natural populations (but see DiBattista, Feldheim, Garant, *et al.*, 2009) and are probably achieved thanks to (i) the initial strong relatedness structure in tested datasets (Csilléry, Johnson, Beraldi, *et al.*, 2006; Gay, Siol & Ronfort, 2013) and (ii) the high versatility of the microsatellite markers developed in the desert locust (i.e., mean expected heterozygosity of 0.84; see also Blondin, Badisco, Pagès, *et al.*, 2013). A drawback of this pedigree-free method is that it is not possible to estimate maternal effects since the identity of mothers are not known. The solution would be to genotype both offspring and parents and to use a parentage assignment method to reconstruct the entire pedigree which could be used afterwards in an animal model either naïve or informed for a maternal effect. Thus, we carried out additional simulations of 100 genotype datasets (still with 16 microsatellite markers), on the crossing scheme that mimicked our experimental design (CS3) and the large optimized crossing scheme (CS15). We showed that the R package **MasterBayes** (Hadfield, Richardson & Burke, 2006) allowed a perfect reconstitution of the original pedigree (i.e. 100% simulated datasets had 0 errors in the reconstructed pedigree). This high performance in pedigree reconstruction is explained by the high levels of information within the Orthopteran microsatellite markers and within the pedigree structures controlled under laboratory conditions (i.e. strong level of relatedness in a half-sib /full-sib design) along with the knowledge of all maternal genotypes (Wang, 2006; Blouin, 2003; Visscher, Hill & Wray, 2008).

### Heritability and maternal effects in phase traits

In order to get first insights into the transmission of phase traits, we measured body color, shape and size traits of late life stages (last-instar larvae and immature adults) of the desert locust under homogeneous conditions of isolation and main other environmental drivers (e.g. humidity, food given *ad libitum*). These measures were acquired under two controlled temperatures, one suboptimal (28°C) and one favoring fast growth (34°C). We used a half-sib/full-sib crossing scheme of 212 individuals maximizing numbers of offspring by family and of dams by sire. Previous studies showed that maternal effects affect the transmission of the F/C ratio, melanization and body weight of hatchlings in *Schistocerca gregaria* (Table 1). The main hypothesis explaining the proximal causes of these maternal effects involves a factor either controlling primary egg size (and thus the amount of yolk) which in turn influences hatchling size and color (Maeno & Tanaka, 2010; Maeno, Piou, Ould Babah, *et al.*, 2013), or released in the egg foam and influencing offspring behavior (Simpson & Miller, 2007).

Despite the statistical limitations of experimental dataset, we combined the simulation results to the experimental results to get some first insights into the transmission of phase traits in the desert locust. First, we showed that the informed model should allow relatively accurate estimates of maternal effect but with low probability (≤ 20%) of detecting a maternal effect of a low or moderate magnitude. Accordingly, we found that no trait exhibits a significant maternal effect (*P*-value ≥ 0.11). Since maternal variances were very low (thus m^2^=0) and additive variance estimates were strictly equal in the the naïve and informed models, we suggest that the transmission of E/F and O/V were not affected by maternal effect. Conversely, a maternal effect might affect body color (m^2^ estimates ~ 0.2) and possibly body size and F/C (m^2^ estimates ~ 0.1). Note that these m^2^ estimates were in all cases (at most twice) lower than the h^2^ estimates from the naïve model.

The relatively low maternal effects estimated from our experimental dataset may be explained by the standardized rearing of the mothers in isolation condition. Doing so, we might both have equalized the maternal environment among our population and remove the main environmental source of maternal effect in the desert locust, i.e. crowding. In addition, maternal effects are expected to be larger for early offspring traits than for late traits (as the ones measured in this study) but can persist into adulthood (McAdam, Garant & Wilson, 2014). In locusts, whether maternal effects detected in hatchlings would persist in later stages is unknown but the colour of the hatchlings changed in the second stadium through the effect of lifetime rearing density from the first stadium (Tanaka & Maeno, 2006). Therefore the maternal variance should be attributed mainly to genetic variation among mothers and to gene-by-environment interaction. For example, the morphometrical and behavioral phases were shown to be transmitted trans-generationally and the genetic variation in this response may indicate a parental effect mediated by parental genes (Chapuis, Estoup, Augé-Sabatier, *et al.*, 2008).

We showed that it is not possible to conclude on heritability estimates with the informed model since power of heritability detection was mostly lower than 5%, whatever the actual heritability of traits. For traits displaying no maternal effect (E/F and O/V), heritability estimates obtained with the naïve model are more reliable even if the power is still limited for heritabilities under 0.3. Therefore E/F and O/V seem to not be (highly) heritable. When maternal effects are present, the naïve model does not allow reliable estimation of heritabilities. Concerning the four traits seemingly affected by maternal effect (green color, brightness, F and F/C), we cannot safely conclude on their level of heritability: the observed changes in heritability estimates between the naive and the informed model could be explained either by a downward bias in h^2^ estimates in the informed model or by an overestimation of h^2^ in the naive model in the presence of maternal effect, as shown by the simulations. Finally, since the maximal larval weight and F/V have heritability and maternal effect estimates in the same order of magnitude, it is also not possible to draw conclusion about their transmission.

Overall, even if our experimental results are not fully conclusive, they might indicate that some phase traits are affected by maternal effects. To increase the probability of formally come to a conclusion on the transmission of phase traits, maternal effects and heritabilities estimates with significantly more power and more accuracy are required. We showed that this may be achieved by optimizing the crossing schemes and more importantly by increasing the sample size. To do so, the use of a pedigree-free method on the available set of microsatellite markers in the desert locust (Blondin et al. 2013), would be promising for future quantitative genetic studies on grouped individuals. Note that this approach requires measuring all traits of interest simultaneously, or at least within the same developmental stadium if individuals are tagged (since tags are lost during molt, Gangwere et al. 1964), before animals are sacrificed for genotyping.

### Conclusion

Our simulations showed that it is challenging to jointly estimate heritability and maternal effects because that it requires datasets with a large sample size and number of families. When it is not possible to get such adequate datasets, conclusions about the heritability of studied traits should remain very cautious and conservative. In any case, comparing the outcomes of both naive and informed models can give precious clues about the impact of maternal effects on heritability assessments. Finally, we want to stress out that 1) simulations are a powerful and convenient tool to explore the performances of potential experimental designs and/or to determine the reliability of obtained estimates and 2) pedigree-free methods may help to achieve satisfying experimental design while limiting the need for time and space.

## Acknowledgements

We thank deeply M. Ould Babah Ebbe, director of the National Anti-Locust Center of Mauritania for providing the egg pods at the start of our experiment. We thank P.-E. Gay for assistance with figures S1 and S2.

